# Human splicing diversity across the Sequence Read Archive

**DOI:** 10.1101/038224

**Authors:** Abhinav Nellore, Andrew E. Jaffe, Jean-Philippe Fortin, José Alquicira-Hernández, Leonardo Collado-Torres, Siruo Wang, Robert A. Phillips, Nishika Karbhari, Kasper D. Hansen, Ben Langmead, Jeffrey T. Leek

## Abstract

We aligned 21,504 publicly available Illumina-sequenced human RNA-seq samples from the Sequence Read Archive (SRA) to the human genome and compared detected exon-exon junctions with junctions in several recent gene annotations. 56,865 junctions (18.6%) found in at least 1,000 samples were not annotated, and their expression associated with tissue type. Newer samples contributed few novel well-supported junctions, with 96.1% of junctions detected in at least 20 reads across samples present in samples before 2013. Junction data is compiled into a resource called `intropolis` available at http://intropolis.rail.bio. We discuss an application of this resource to cancer involving a recently validated isoform of the ALK gene.

## 1 Preliminaries

Gene annotations such as those compiled by RefSeq [1] and GENCODE [2] are derived primarily from alignments of spliced cDNA sequences and protein sequences [3,4]. So far, the impact of RNA sequencing (RNA-seq) data on annotation has been limited to a few projects including ENCODE [5] and Illumina Body Map 2.0 [6].

To measure how much splicing variation present in publicly available RNA-seq datasets is missed by annotation, we aligned 21,504 Illumina-sequenced human RNA-seq samples from the Sequence Read Archive (SRA) to the *hg19* genome assembly with Rail-RNA [7] and compared exon-exon junction calls to exon-exon junctions from annotated transcripts. We compared exon-exon junctions rather than full transcripts because junction calls from short RNA-seq reads are considerably more reliable than assembled transcripts [8]. Details of our alignment protocol are reviewed in Methods. All alignment was performed in the cloud using Amazon Web Services (AWS) Elastic MapReduce, costing 72 US cents per sample, as computed in Methods. We considered only Illumina platforms because of their ubiquity and high base-calling accuracy. Specifically, the samples we aligned were obtained by querying the SRA metadata SQLite database of the R/Bioconductor package SRAdb [9] as of April 2015 for all Illumina-sequenced human RNA-seq samples. In the remainder of this paper, we use the term “annotation” to refer to junctions from the union of transcripts across several gene annotation tracks from the UCSC Genome Browser [10]. For *hg38* annotations, coordinates were lifted over to *hg19.* See Methods for details and Table S1 for included gene annotations together with the number of junctions in each. In all, we found 542,706 annotated junctions: 506,105 were present in annotations of *hg19,* and the rest were added by annotations of *hg38.*

## 2 Results

We compiled the junction calls and associated coverage levels for 21,504 SRA RNA-seq samples into a resource called `intropolis` available at http://intropolis.rail.bio. Using this resource, we addressed several questions that are fundamental to our understanding of the transcriptome and informative for analyses by individual investigators.

### 2.1 Robustness

We first asked whether our junction calls were robust across alignment protocols. The SEQC/MACQ-III consortium (hereafter called SEQC) aligned a subset of 1,720 universal human reference RNA and human brain reference RNA samples [11] of the 21,504 samples we considered using three different protocols: NCBI Magic [12], r-make (which uses STAR [13]) and Subread [14]. Junctions called by Rail-RNA are compared with junctions called by SEQC across the subset in Figure 1. Of junctions found by Rail-RNA in at least 80 SEQC samples, as many as 97.5% are found by at least one SEQC alignment protocol, and 90.1% are found by all three. Note that 80 SEQC samples is 4.7% of 1,720, comparable to a 1,000-sample threshold discussed below for the 21,504 SRA samples.

**Figure 1.**
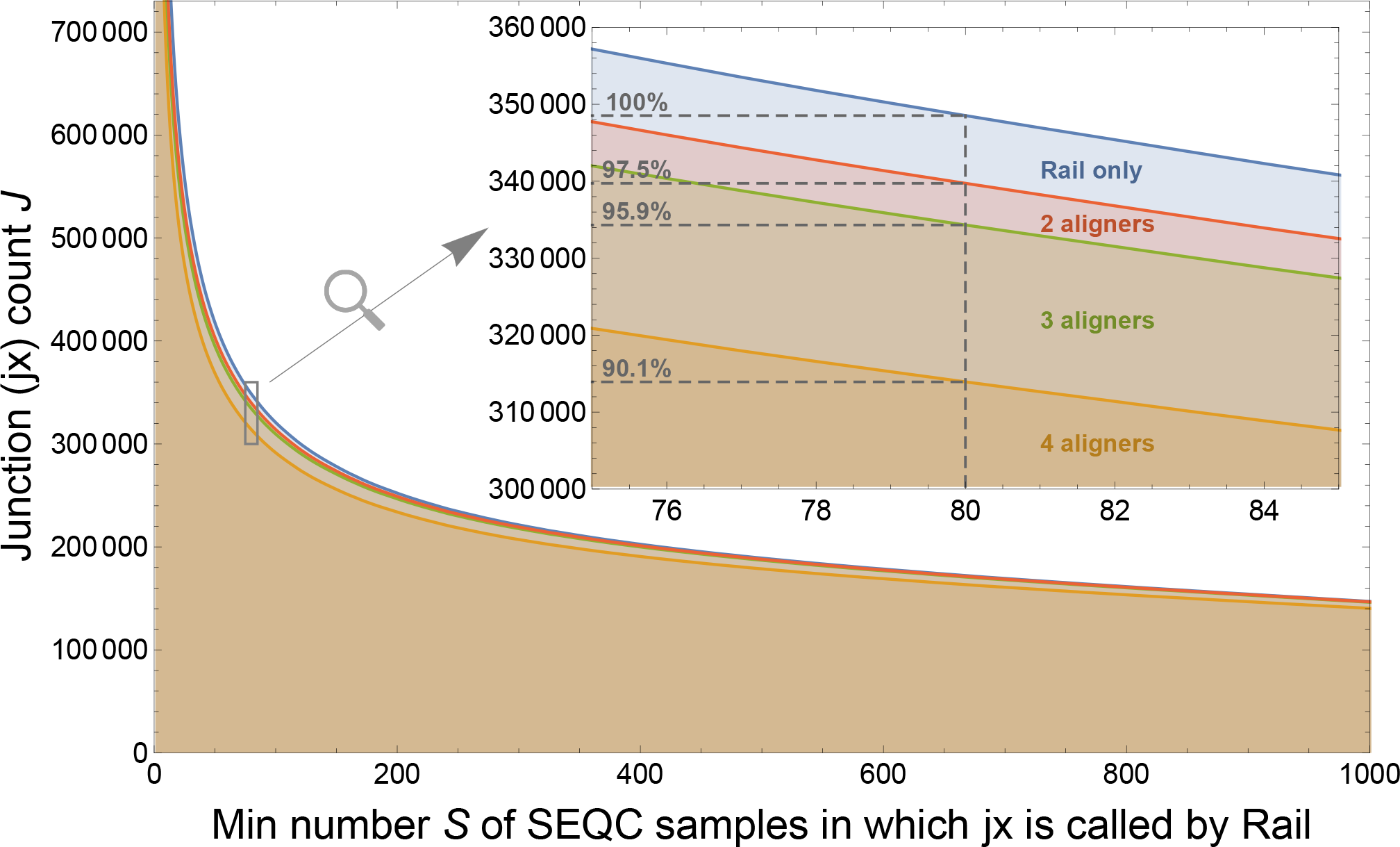
Displayed is the number of exon-exon junctions J found by Rail-RNA and other alignment protocols in at least S of the 1,720 brain and universal human reference RNA-seq samples also studied by the SEQC/MACQ-III consortium [11] (i.e., SEQC). “2 aligners” (red), “3 aligners” (green), and “4 aligners” (orange) refer to junctions we found with Rail-RNA that were also found by, respectively, 1, 2, and 3 of the alignment protocols used by SEQC. 97.5% of junctions found by Rail-RNA in at least 80 SEQC samples were also found by another SEQC alignment protocol, and 90.5% were also found by all three SEQC alignment protocols.

### 2.2 Relationship between annotation and expression of splice junctions

We next asked whether annotated junctions represent the diversity of junction expression observed in the population at large. We considered a junction well-supported in our data if it appeared in a large number of samples. We calculated the number of junctions that appeared in at least S samples across a range of cutoffs. For each junction we considered, we also evaluated whether it appeared in annotation. We considered the following levels of evidence: (1) fully annotated junctions; (2) separately annotated junctions (typically exon skipping events), where both the donor and acceptor sites appear in one or more junctions from annotation, but never for the same junction; (3) alternative donor and acceptor sites, where only either the donor or the acceptor site appears in one or more junctions from annotation; and (4) novel junctions, where neither donor nor acceptor site is found in any annotated junction.

We observed that the junctions most widely expressed across samples and experiments were well-documented in annotation. For example, we observed that junctions that appeared in at least 40% of human RNA-seq samples on SRA (*S ≥* 8,000) were also present in previous annotation at least 99.8% of the time. However, 18.6% of junctions that appeared in 1,000 or more samples did not appear in annotation (Figure 2a). Many of these unannotated junctions are partially annotated, but 3.5% of junctions found in over 1,000 samples do not match any splice site from an annotated junction.

**Figure 2.**
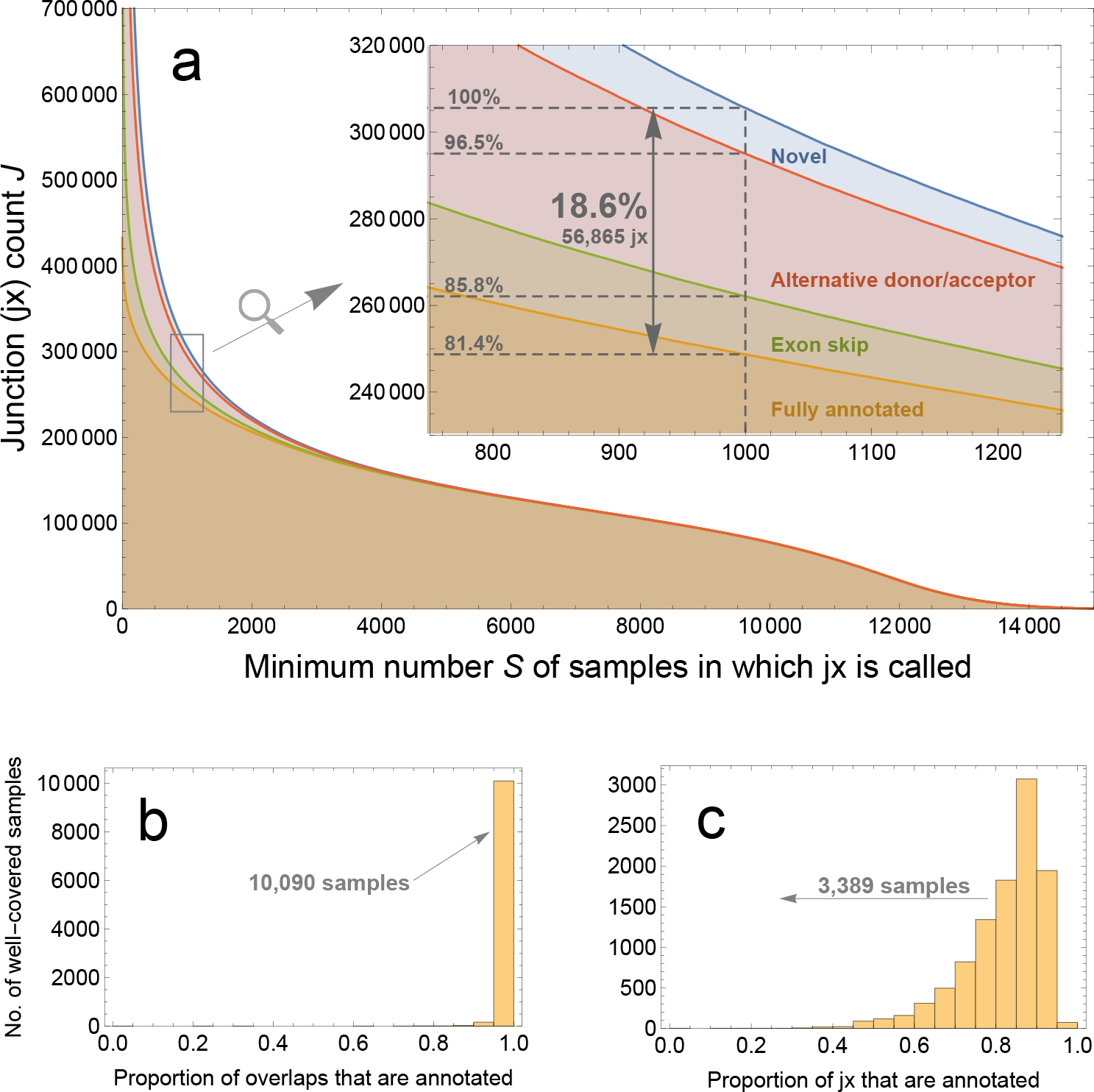
(a) shows how many exon-exon junctions J are found in at least S samples of the 21,504 human RNA-seq samples on SRA aligned here. It also shows how much evidence for these junctions is found in gene annotation: “fully annotated” (orange) means the junction is in an annotated transcript, “exon skip” (green) means a called junction’s donor and acceptor sites are annotated in distinct junctions, “alternative donor/acceptor” (red) means only one of a called junction’s donor and acceptor sites is in a junction from annotation, and “novel” (blue) means neither donor nor acceptor site is annotated. The overwhelming majority (99.7%) of junctions found in over 8,000 samples are fully annotated, but 18.6% found in over 1,000 samples are not fully annotated. (b) and (c) restrict attention to the 10,311 samples for which 100,000 junctions are discovered in each. (b) refers to overlaps as defined in the main text, and it shows that if a sample’s junctions are weighted by the number of reads that map across them, annotation captures over 95% of variation in 10,090 samples. But if each junction is weighted equally as in (c), annotation captures less than 80% of variation in 3,389 samples.

We also took an investigator-focused view of the relationship between annotation and expression. Most investigators collect only a small number of samples for their study. We restricted attention to samples where at least 100,000 junctions were found to rule out obviously small RNA-seq samples and samples that were mislabeled as RNA-seq on SRA. In each sample, we counted the number of instances where a read maps across a junction. (A read mapping across two junctions thus contributes two instances.) The total number of such “junction overlaps” across samples is a measure of the total expression of junctions across SRA. Most of the reads that map to junctions map to annotated junctions (Figure 2b). In 10,090 of a total of 10,311 samples that meet our criterion of 100,000 junctions observed, over 95% of junction overlaps correspond to annotated junctions.

This represents only the bulk coverage of junctions. We can also consider the number of junctions observed, regardless of coverage. In 3,389 out of 10,311 samples, we observe that fewer than 80% of junctions appear in annotation (Figure 2c). So while the most highly covered junctions are well-annotated, there is a large number of junctions that are well-covered across multiple samples but may not appear in any given small subset of samples.

To explore this idea further, we investigated the potential for single studies to be the sole contributors of individual unannotated junctions. In this event, the junction may not have been called robustly across experimental protocols. Here, we considered junctions that appeared in at least P projects instead of samples. We again broke this calculation down by the different potential levels of evidence: whether the junction was entirely novel, had an alternative donor or acceptor, an exon skip, or whether it was fully annotated (Figure 3). The story at the project level mirrors the story at the sample level: 23.4% of junctions found in over 200 of the 929 projects are not fully annotated. So unannotated junctions recur across independent investigations.

**Figure 3.**
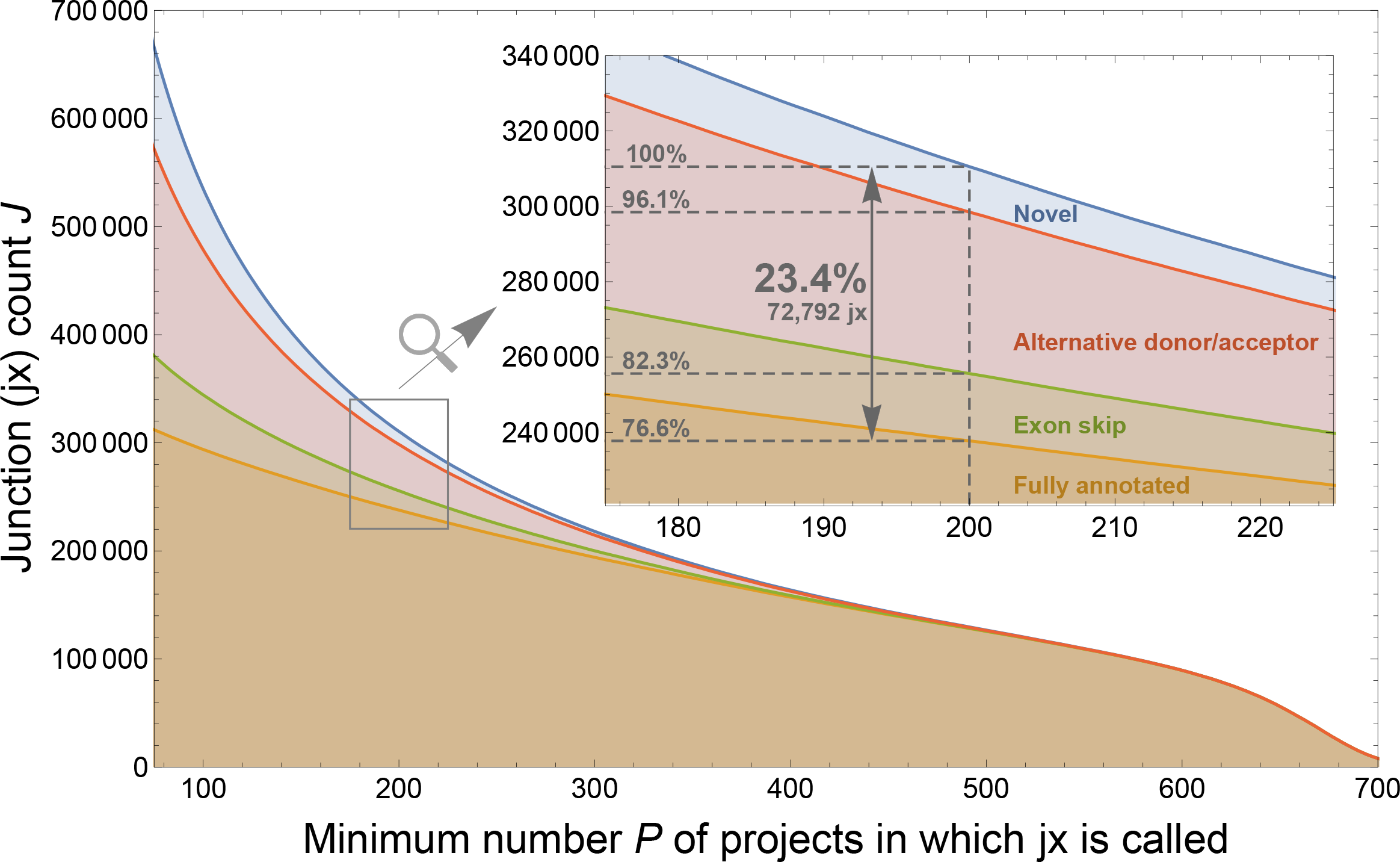
Displayed is the number of exon-exon junctions J found in at least P projects of the 929 human RNA-seq projects on SRA considered in this paper. It also shows how much evidence for these junctions is found in gene annotation: “fully annotated” (orange) means the junction is in an annotated transcript, “exon skip” (green) means a called junction’s donor and acceptor sites are annotated in distinct junctions, “alternative donor/acceptor” (red) means only one of a called junction’s donor and acceptor sites is in a junction from annotation, and “novel” (blue) means neither donor nor acceptor site is annotated. 23.4% of junctions found in over 200 projects are not fully annotated.

### 2.3 Technical and biological variation in junction expression across samples

We next explored variation across the 21,504 samples we processed. We wanted to see the combination of technical and biological factors that contribute to variation in unannotated junction expression. In this analysis, we considered only the 56,865 unannotated junctions found in at least 1,000 samples of the 21,504, and the subset of 21,057 samples of the 21,504 with at least 100,000 reads each. We performed a Principal Component Analysis (PCA) on the data matrix where rows correspond to the 56,865 unannotated junctions and columns correspond to the 21,057 samples. (See Methods for technical details of the decomposition.)

PC1 explains the overwhelming majority of the variance (87.9%) and has a Pearson correlation coefficient *r* = 0.978 with junction sequencing depth *s_j_* as measured by total junction overlaps (i.e., instances where a read maps across a junction) in sample *j* (Figure 4) after normalization by library size and log transformation. PC1 is also highly correlated with log-transformed read length *l_j_* (*r* = 0.639), but much less correlated with log-transformed total number of mapped reads *C_j_* (*r* = 0.277), showing that enrichment for splice junctions is different in different samples. (See Methods for precise definitions of correlates.)

**Figure 4.**
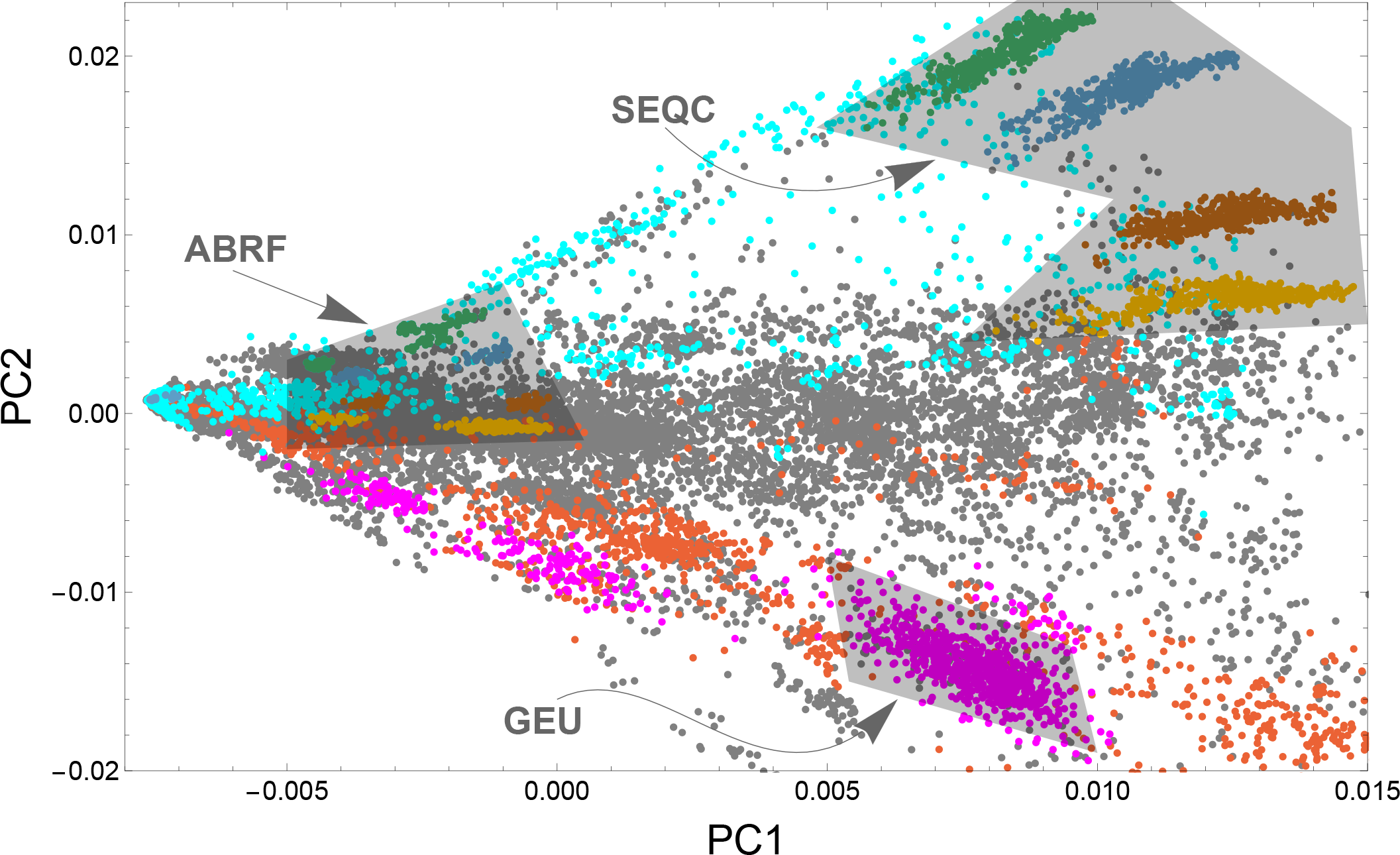
Displayed is the first principal component (PC1) vs. the second principal component (PC2) for a Principal Component Analysis (PCA) with a coverage data matrix where rows are junctions and columns are samples. (See Methods for technical details.) Each point corresponds to a distinct sample. Gray points are unlabeled samples, red points are blood samples, magenta points are lymphoblastoid cell line samples, and cyan points are brain samples. GEUVADIS (GEU) is a sizable cluster of magenta points. The ABRF and SEQC consortia each sequenced mixtures of Universal Human Reference RNA (UHRR) and Human Brain Reference RNA (HBRR) in four sample ratios UHRR:HBRR that form distinct clusters in the shaded regions: 0:1 (green), 1:3 (blue), 3:1 (brown), and 1:0 (yellow).

We further examined samples belonging to specific groups that generated well-characterized datasets. Both the SEQC consortium and ABRF [15] studied universal human reference RNA (UHRR) and human brain RNA reference (HBRR) samples constructed by the MACQ-III consortium for quality control. UHRR is comprised of total RNA from 10 different cancer cell lines representing various human tissues, while HBRR samples are comprised of total RNA from several donors across several brain regions. Both groups studied these samples in four different mixture ratios—0:1, 1:3, 3:1, and 1:0—with each sample sequenced at multiple sites. The four mixtures separate well, and each lies on a radial line passing through the singular point on the left. Data from the two groups are separated because they used different sequencing depths and read lengths.

The four SEQC UHRR:HBRR sample ratios form four clusters distinguished by PC2, and the ABRF UHRR:HBRR sample ratios form clusters distinguished by both PC1 and PC2. Observe that there is a singular point where all points appear to converge (Figure 4). Here, the number of junctions detected in a sample approaches zero. A radial line extending from the singular point rotating clockwise across the plot passes over UHRR:HBRR sample ratios in the same order for ABRF as it does for SEQC. Though ABRF and SEQC have some overlap in managing investigators, they are two different projects that employed randomized study designs, making a strong case that PC2 is distinguishing mostly biological rather than technical factors.

Lymphoblastoid cell lines, typically made from HapMap samples, are extensively present in SRA. Different studies cluster together and are again placed on a radial line going through the singular point; each study used very different sequencing depths and read lengths. Searching the SRA metadata, we could classify a number of samples as brain and blood. Again, these samples fall along radial lines through the singular point. The biggest separation in PC2 is between brain and blood, two tissue types that are well-represented in SRA.

### 2.4 Novel junction discovery over time

We proceeded to measure the accumulation of “confidently called” junctions over calendar time. A junction was “confidently called” if it was found in at least 20 reads across all samples. We measured the discovery date of a junction as the earliest submission date to the BioSample database [16] from among all samples in which the junction was found by Rail-RNA. The ≥ 20-read curve has noticeable spikes in 2009 and 2011 but appears to decelerate significantly before 2013, by which time 96.1% of junctions were discovered.

Recent samples added to SRA have contributed few novel junctions. Curves for more stringent coverage thresholds (Figure 5) level off sooner; the curve for the most stringent threshold (≥ 160 reads) is essentially flat by 2012. Ranked and labeled are the dominant contributing projects from days on which the most junctions were discovered. The largest single contribution comes from UWE, the University of Washington’s Human Reference Epigenome Mapping Project [17], on April 4, 2011, when 252,628 new junctions appeared. The submission includes total RNA from fetal tissue, which exhibits markedly different expression than adult tissue [18]. The second, third, fourth, and fifth-largest contributions are from, respectively, ENCODE [19], early studies of 69 lymphoblastoid cell lines (LCLs) [20] and 41 Coriell cell lines [21], and the Illumina Body Map 2.0 sequencing of 16 human tissue types [6]. The GEUVADIS submission of 464 LCLs is on only the 55th-largest contributing date, November 7, 2012. By this time, LCLs had already been well-studied using RNA-seq.

**Figure 5.**
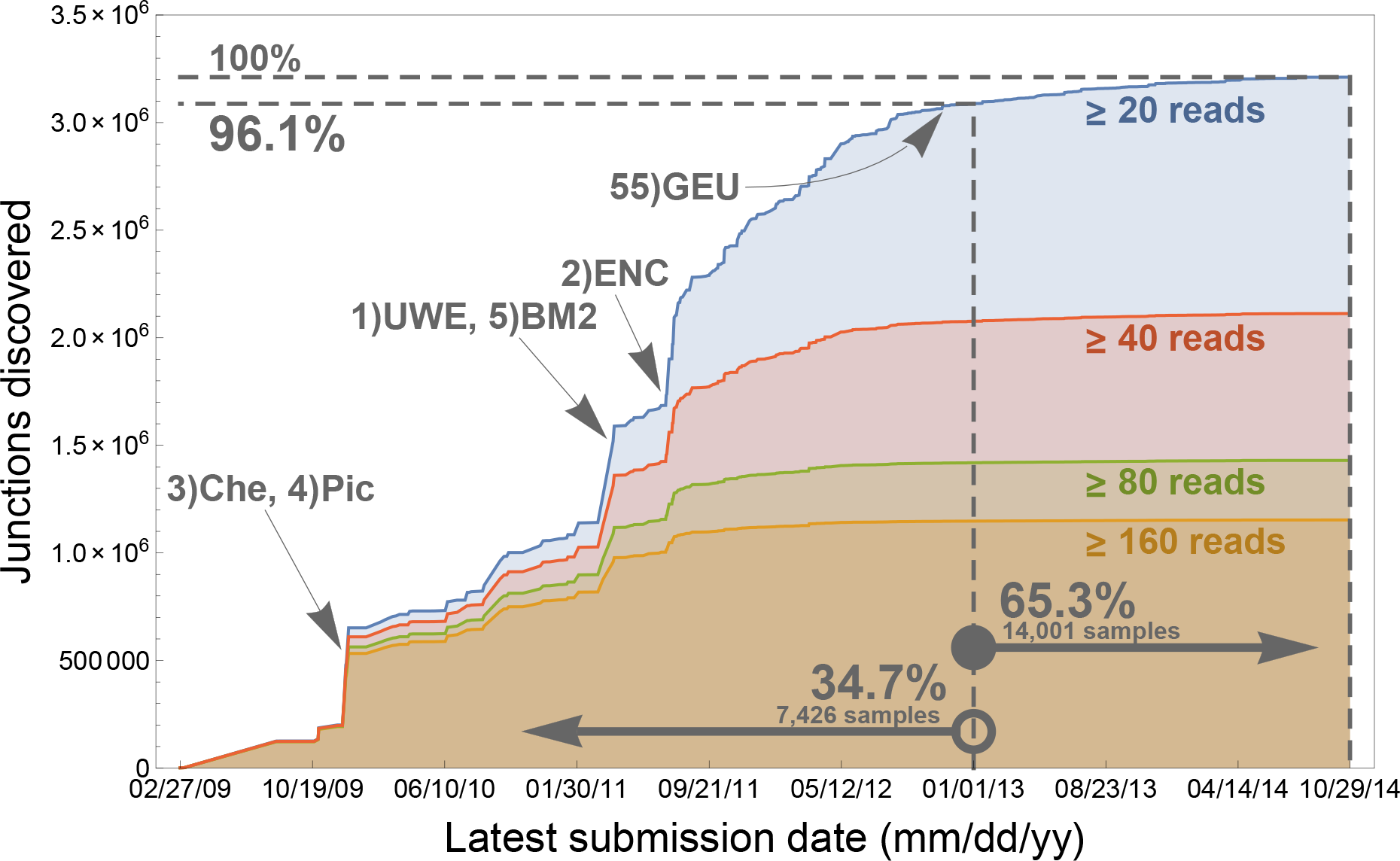
The 3,211,228 junctions found in at least 20 reads across samples are accumulated by their “discovery dates.” Here, discovery date of a junction is taken to be the earliest submission date to the BioSample database from among the samples in which the junction was found. 96.1% of the junctions were discovered before January 1, 2013, despite how only 34.7% of samples depicted in the figure had been submitted by then, and afterwards discovery levels off. Demanding higher levels of confidence (the red, green, and orange curves) gives rise to earlier asymptotes. Ranked from 1 to 5 are the dominant contributing projects from dates on which the most junctions are discovered. “Che” refers to a study of 41 Coriell cell lines by Cheung et al. [21], “Pic” refers to a study of 69 LCLs by Pickrell et al. [20], “UWE” refers to the University of Washington Human Reference Epigenome Mapping Project [17], “BM2” refers to Illumina Body Map 2.0 [6], and “ENC” refers to ENCODE [19]. “GEU” refers to GEUVADIS [32], whose 464 LCLs uncovered few junctions that had not already been discovered.

To determine whether the annotation of junctions is being driven by RNA-seq experiments, we examined the correlation between annotated junctions and the discovery date of observed junctions over calendar time. GENCODE released 18 versions between September 2009 and December 2012. Call a confidently called junction “documented” if it appears in at least one GENCODE release. Most documented junctions (80.0%) appear in the earliest GENCODE release (Figure 6a). Documented junctions tend to have early discovery dates (Figure 6b); in fact, by late January 2010, 74.2% of documented junctions were discovered, while 20.3% of confidently called junctions were discovered (Figure 6c). This makes sense: annotated junctions tend to be found in many samples, making it likelier at least one sample has an early submission date to BioSample. It is reasonable to speculate that there is a correlation between junction discovery date and GENCODE appearance date: perhaps shortly after a junction is discovered, it appears in GENCODE. But inspection of the relationship between documentation date and discovery date suggests that only the first GENCODE release introduced new junctions with significantly earlier discovery dates than other releases (Figure 6b). The reason for this phenomenon is junctions appearing first in GENCODE’s first release are present in many more samples (median = 5,817) than junctions appearing first in other GENCODE releases (median = 603 samples) (Figure 6d).

**Figure 6.**
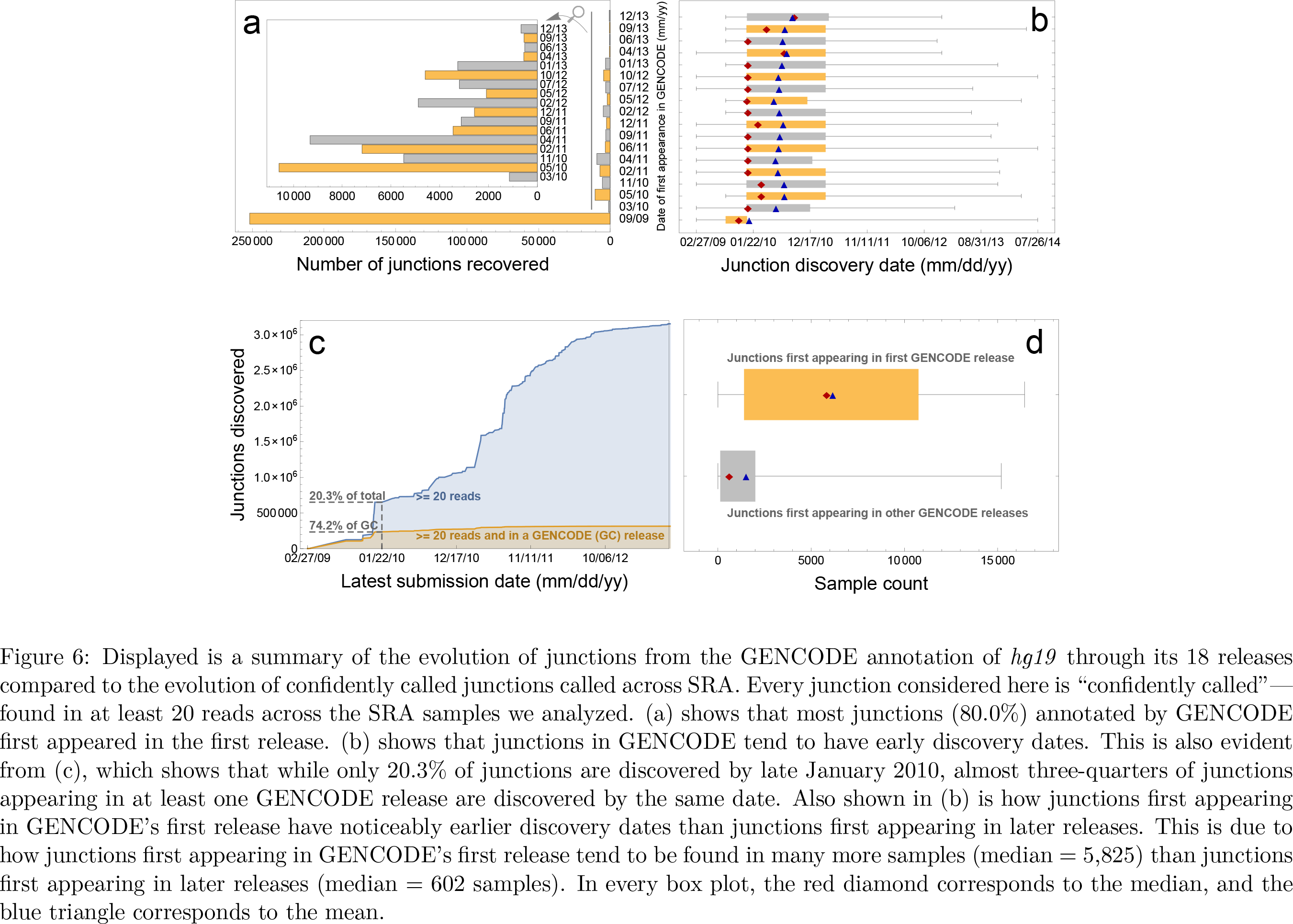
Displayed is a summary of the evolution of junctions from the GENCODE annotation of *hg19* through its 18 releases compared to the evolution of confidently called junctions called across SRA. Every junction considered here is “confidently called”— found in at least 20 reads across the SRA samples we analyzed. (a) shows that most junctions (80.0%) annotated by GENCODE first appeared in the first release. (b) shows that junctions in GENCODE tend to have early discovery dates. This is also evident from (c), which shows that while only 20.3% of junctions are discovered by late January 2010, almost three-quarters of junctions appearing in at least one GENCODE release are discovered by the same date. Also shown in (b) is how junctions first appearing in GENCODE’s first release have noticeably earlier discovery dates than junctions first appearing in later releases. This is due to how junctions first appearing in GENCODE’s first release tend to be found in many more samples (median = 5,825) than junctions first appearing in later releases (median = 602 samples). In every box plot, the red diamond corresponds to the median, and the blue triangle corresponds to the mean.

## 3 Application to *ALK* isoform discovery

We have compared the variation in our database intropolis to standard gene annotations. intropolis associates each junction with the set of samples where the junction was called and the number of reads spanning the junction in that sample, enabling biological investigators to gain new insights. Here, we give a simple example application involving the anaplastic lymphoma kinase (*ALK*) gene.

*ALK* is frequently mutated or aberrantly expressed in cancers including neuroblastoma [22–25] and non-small-cell lung adenocarcinoma, where in particular it has been found to participate in the fusion gene *EML4-ALK* [26]. Cancers with *ALK* abnormalities are often responsive to treatment with *ALK* inhibitors such as crizotinib [27]. *ALK* is a good therapeutic target because it is rarely expressed in normal adult tissue [28]. A novel ALK transcript variant present in about 11% of melanomas and occasionally in other cancer subtypes was recently identified [29]. The transcript is described as resulting from a *de novo* alternative transcription initiation (ATI) site in *ALK* intron 19 and is dubbed *ALK^ATI^*. The kinase activity of *ALK^ATI^* is found to be suppressed by various ALK inhibitors, and a patient with *ALK^ATI^*-expressing metastatic melanoma is shown to exhibit significant tumor shrinkage after treatment with crizotinib.

To investigate the prevalence of *ALK^ATI^* on SRA, we searched for a deficit of junction expression in *ALK* exons 1−19 compared to exons 20−29. We did this by defining a junction inclusion ratio D measuring to what degree junctions between exons 20-29 are expressed relative to junctions between exons 1-19 (see Methods). This signature is a necessary but not sufficient condition for exclusive *ALK^ATI^* expression: the expression signature also arises in, for example, the *EML4-ALK* fusion gene. Table S2 shows the ten top SRA samples we studied ranked in order of decreasing D. As expected, four such samples are cancers, including uveal melanoma. Three of the ten samples are from two melanocyte cell cultures studied as part of the ENCODE project, “NHEM_M2” and “NHEM.LM2.” Cap analysis of gene expression (CAGE) data from ENCODE on the same cell lines shows a TSS within *ALK* intron 19, where the TSS was localized for *ALK^ATI^* (Supplementary Fig. S1). This raises the possibility that the transcript is expressed in normal melanocytes. While Wiesner et al. found no *ALK^ATI^* expression in 1,600 samples from 43 different normal tissues across the GTEx project, including skin, it should be noted that melanocytes comprise only up to 10% of skin cells. In addition to melanocytes, the *ALK^ATI^* transcript may be expressed in macrophages. We also observed that the macrophage and macrophage+fibroblast samples from Table S2 are part of the study [30] that additionally sequenced the same samples exposed to tumor necrosis factor (TNF). The two samples exposed to TNF appear to have no expression of the *ALK* gene, suggesting that TNF may participate in suppressing *ALK* gene expression. This is supported by [31] in lymphoma.

## 4 Discussion

We have measured variation in junction expression across thousands of RNA sequencing samples. Our analysis demonstrates both the strengths and weaknesses of relying on current annotation for RNA-seq analysis. We have also used our population-level view of transcription to understand the potential hazards of analyzing individual samples without a clear understanding of the background variation in junction discovery levels. We have introduced a resource, `intropolis`, for others to investigate junction variation, and we have provided an example of the utility of our resource in the case of *ALK* gene expression.

As highlighted by Figure 2a-b, considering only the variation contained in annotation may suffice if an investigator is interested only in well-expressed transcript isoforms. However, genes that are not generally well-expressed and nonetheless present in a small but significant number of samples on SRA are likelier to be incompletely annotated. Our approach to synthesizing large public RNA-sequencing datasets offers the opportunity to study transcription more deeply than ever before.

## Data availability

`intropolis` may be downloaded at http://intropolis.rail.bio.

## Acknowledgments

AN, LCT, SW, KDH, BL, and JTL were supported by NIH/NIGMS grant 1R01GM105705 to JTL. AN was supported by a seed grant from the Institute for Data Intensive Engineering and Science (IDIES) at Johns Hopkins University to BL and JTL. LCT was supported by Consejo Nacional de Ciencia y Tecnología Mexico 351535. BL was supported by a Sloan Research Fellowship to BL. JAH was supported by Fundación UNAM. Alignment using Amazon Web Services was partially supported by an AWS in Education grant to BL. SW, RAP, and NK were supported through the Johns Hopkins Center for Computational Biology summer internship program.

## Methods

### Identifying annotated junctions

Following [33], we extracted junctions from transcripts across all the latest “empirical” gene annotation tracks in the UCSC Genome Browser [10] for *hg19* and *hg38* except GENCODE [2] and Ensembl [34]. (While GENCODE’s tracks are also in the UCSC Genome Browser, we chose to download them from the GENCODE website http://www.gencodegenes.org/releases/instead: as of January 24, 2016, GENCODE v22 was the latest GENCODE track listed, but GENCODE v24 had already been released.) Empirical tracks are based on alignments of e.g. spliced cDNA and protein sequences and are listed in Table S1. Annotation tracks based on algorithmic predictions from genome sequence (Augustus, GeneID, Genscan, N-SCAN, and SGP) were excluded because they are comprised of transcripts that were not directly observed in experiment. Ensembl was excluded because GENCODE is already a merge of Ensembl and HAVANA transcripts. After junction coordinates were extracted, all *hg38* coordinates were lifted over to *hg19* where feasible, and the union of all junctions was taken. Since the `intropolis` database was formed from alignments to only the *hg19* chromosomal assembly, only those junctions corresponding to the *hg19* chromosomal assembly were kept to form a final list of annotated junctions. Table S1 lists all gene annotations used to determine our set of annotated junctions. We froze these annotations on January 24, 2016 and compressed them into an archive available at http://verve.webfactional.com/misc/jan_24_2016_annotations.tar.gz. We ran the script https://github.com/nellore/runs/blob/master/sra/rip_annotated_junctions.pywithPyPyv2.5.0 to extract junctions from these annotations, performing coordinate conversions from *hg38* to *hg19* where appropriate. The final list of junctions we defined as “annotated” is available at https://github.com/nellore/runs/blob/master/sra/annotated_junctions.tsv.gz.

### Selecting SRA samples

Samples were selected by querying the SRA metadata SQLite database of the R/Bioconductor package SRAdb [9]. The database was downloaded from http://gbnci.abcc.ncifcrf.gov/backup/SRAmetadb.sqlite.gz, but this file is updated regularly. The version of SRAmetadb.sqlite.gz we used was updated April 1, 2015, and it is available at ftp://ftp.ccb.jhu.edu/pub/langmead/sra_junctions/SRAmetadb.sqlite.gz. We selected all `run_accessions` from the `sra` table with `platform =’ILLUMINA’, library.strategy =’RNA-Seq’`, and `taxon_id = 9606` (human) that also had URLs for FASTQs on the European Bioinformatics Institute server listed in the `fastq` table. Our query may be reproduced with the script https://github.com/nellore/runs/blob/master/sra/define_and_get_fields_SRA.R compatible with R v3.1.0.

### Alignment with Rail-RNA

Rail-RNA v0.1.7b [7] was used for alignment. We aligned to *hg19* rather than the more recent *hg38* assembly because of *hg19*’s continued prevalence, including use by the GEUVADIS consortium [32] in its study of 462 lymphoblastoid cell line (LCL) samples as well as the GTEx consortium [35] in its ongoing large-scale study of gene expression across human tissues. We performed a single pass of alignment; that is, reads were not realigned after junctions were discovered to improve alignments of short-anchored reads. Alignment was performed in the cloud using AWS Elastic MapReduce on Elastic Compute Cloud spot instances, i.e., standardized units of computing capacity. Spot instances permit bidding for compute to save money, where bids that equal or exceed a market price are fulfilled. However, if the market price drops below a bid, instances could be lost, and a computational job could fail. So saving money by bidding for spot instances comes with risk, and rather than aligning all samples in one batch, we distributed this risk by dividing alignment up into 43 batches of about 500 samples each. Analysis of each batch was itself divided into (1) a preprocessing job flow, which downloaded and preprocessed compressed FASTQs from the European Bioinformatics Institute’s mirror of SRA, writing results to Amazon’s cloud storage service S3; and (2) an alignment job flow, which was configured to write only exon-exon junction coordinates and the number of reads in each sample mapping across each detected junction. Each preprocessing job flow was run on a cluster of 21 c3.2xlarge instances, each with 8 Intel Xeon E5-2680 v2 (Ivy Bridge) processing cores and 15 GB of RAM. Each alignment job flow was run on a cluster of 61 c3.8xlarge instances, 32 Intel Xeon E5-2680 v2 (Ivy Bridge) processing cores and 60 GB of RAM. Summing the sizes of the 43 compressed files output by the 43 runs gives 5.3 GB, about the size of an alignment BAM for a single RNA-seq sample. Our alignment runs may be reproduced by following the instructions at https://github.com/nellore/runs/blob/master/sra/README.md.

### Alignment cost

Alignment was performed over a period of eight days. 21,506 samples spanning 62.2 trillion nucleotides were initially selected for alignment, but two samples (run accession numbers SRR651690 and DRR023700) were not found on the European Bioinformatics Institute server and were therefore excluded. We used the Amazon Cost Explorer to compute total cost; summing across eight days of activity, it came to US$15,393.69, or 72 cents per sample. Costs divided up by Amazon service over the period of computational activity may be viewed at https://github.com/nellore/runs/blob/master/sra/hg19.costs.csv.

### Reproducing main figures

All data underlying Figures 1, 2, 3, 5, and 6 are reproducible with the Python v2.7 script https://github.com/nellore/runs/blob/master/sra/tables.py, which was run using PyPy v2.5.0. These figures as well as Figure 4 were generated with the Mathematica v10.3.1 notebook https://github.com/nellore/runs/blob/master/sra/preprint_figures.nb. SEQC/MAQC-III consortium junction data was downloaded from http://www.nature.com/nbt/journal/v32/n9/extref/nbt.2957-S4.zip. BioSample submission dates for 77 SRA runs (0.3% of the samples we studied) were not found on the server, so these runs were excluded from the analyses involving junction discovery dates presented in Figures 5 and 6.

### Analysis of novel *ALK* isoform

The junction inclusion ratio D discussed in the main text is defined as follows. Suppose the number of instances where junctions are overlapped by reads (i.e., the junction overlap count) in *ALK* exons 1-19 is *A,* and the junction overlap count in *ALK* exons 20-29 is *B.* The normalized difference *D* = (*B* – *A*)/(*A* + *B*) is close to 1 when exons 1-19 are unexpressed compared to exons 20-29, and close to −1 when exons 20-29 are unexpressed compared to exons 1-19.

The *ALK* analysis may be reproduced by first filtering `intropolis` for junctions in ALK with the script https://github.com/nellore/runs/blob/master/sra/alk.sh, and then running the Mathematica 10.3.1 notebook https://github.com/nellore/runs/blob/master/sra/alk.nb. Samples found were checked manually for their descriptions on SRA at http://www.ncbi.nlm.nih.gov/sra, and the UCSC Genome Browser screenshot of Figure S1 was created using the Genome Browser’s PDF/PS utility.

### Principal component analysis

Restrict attention to unannotated junctions found in at least 1.0 of the 21,504 SRA samples we studied and further to only those samples with at least 100.0 reads each. Consider the number of reads C_j_ overlapping the ith unannotated junction in the *j*th sample. We formed the normalized log-counts 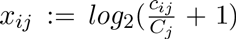, where *C_j_* is the number of mapped reads for sample *j*. We then used the row-centered matrix A for PCA; that is, *A_ij_* = 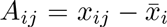. More specifically, we computed the cross-product *A_t_A* in a block-wise manner, and we subsequently performed a singular value decomposition (SVD) of A* A to obtain the right-singular vectors (principal components) with a randomized SVD algorithm [36]. Three correlates of PC1 are mentioned in the text. They are defined as

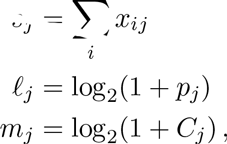

where *j* indexes samples and *p_j_* is the read length in sample *j*.

Scripts for reproducing the PCA analysis are available in the **sra** subdirectory of https://github.com/nellore/runs and described in https://github.com/nellore/runs/blob/master/sra/README.md. Output of the analysis sourced the Mathematica 10.3.1 notebook https://github.com/nellore/runs/blob/master/sra/preprint_figures.nb for generating Figure 4.

## Supplementary Figures and Tables

**Table S1.** Gene annotations from which exon-exon junctions were extracted and unioned to obtain a list of annotated junctions. All tracks were taken from the UCSC Genome Browser [10] except for GENCODE [2], which was downloaded from the GENCODE website http://www.gencodegenes.org/releases/. Junction coordinates from *hg38* annotations were lifted over to *hg19* before the union was performed. Of all gene annotations listed here, the Swedish Bioinformatics Institute (SIB) genes has the most, with over 400,000 junctions for each of *hg19* and *hg38.*

| Gene annotation | Number of junctions | Reference |
| --- | --- | --- |
| *hg19* |  |  |
| AceView | 395,410 | [12] |
| CCDS | 174,337 | [37] |
| GENCODE v19 | 343,887 | [2] |
| UCSC | 264,339 | [10] |
| lincRNAs | 30,256 | [6] |
| MGC | 160,388 | [38] |
| RefSeq | 235,512 | [1] |
| SIB | 414,422 | [39] |
| Vega | 242,672 | [2] |
| *hg38* |  |  |
| CCDS | 178,775 | [37] |
| GENCODE v24 | 346,547 | [2] |
| UCSC | 356,644 | [10] |
| lincRNAs | 29,940 | [6] |
| MGC | 169,302 | [38] |
| RefSeq | 247,577 | [1] |
| SIB | 433,606 | [39] |

**Table S2.** Top ten samples across the 21,504 analyzed in this paper in order of descending junction inclusion ratio *D,* as defined in the table. D essentially measures the difference in expression between junctions across ALK exons 1-19 and junctions across *ALK* exons 20-29. Values of D close to 1 may point toward expression of *ALK^ATI^*, a novel transcript variant of ALK recently identified in [29] across several cancers but not normal cells. Several cancer samples appear, but interestingly, normal cell samples also appear, including melanocytes and macrophages.

| Rank | Sample (i.e., run) | Project | Description of sample | Junction coverage A for ALK exons 1-19 | Junction coverage B for ALK exons 20-29 | Total junction coverage C across ALK | $D=(B-A)/C$ |
| --- | --- | --- | --- | --- | --- | --- | --- |
| 1 | SRR545713 | SRP007461 | NHEM.f_M2: normal human melanocyte | 0 | 139 | 139 | 1 |
| 1 | SRR396804 | SRP010166 | non-small cell lung adenocarcinoma | 0 | 172 | 172 | 1 |
| 1 | SRR620100 | SRP017262 | leukemia | 0 | 108 | 108 | 1 |
| 4 | SRR1289650 | SRP042031 | macrophage | 1 | 85 | 86 | 0.976 |
| 5 | SRR1289651 | SRP042031 | macrophage cultured with fibroblast | 1 | 77 | 78 | 0.974 |
| 6 | SRR545716 | SRP007461 | NHEM_M2: normal human melanocyte | 2 | 94 | 96 | 0.958 |
| 7 | SRR628586 | SRP017413 | uveal melanoma | 12 | 111 | 123 | 0.805 |
| 8 | DRR016705 | DRP001919 | H2228, an EML4-ALK-expressing lung adenocarcinoma cell line | 38 | 285 | 333 | 0.765 |
| 9 | SRR545714 | SRP007461 | NHEM.f_M2: normal human melanocyte | 14 | 63 | 77 | 0.636 |
| 10 | ERR532612 | ERP006077 | prostate tumor | 16 | 53 | 69 | 0.536 |

**Figure S1.**
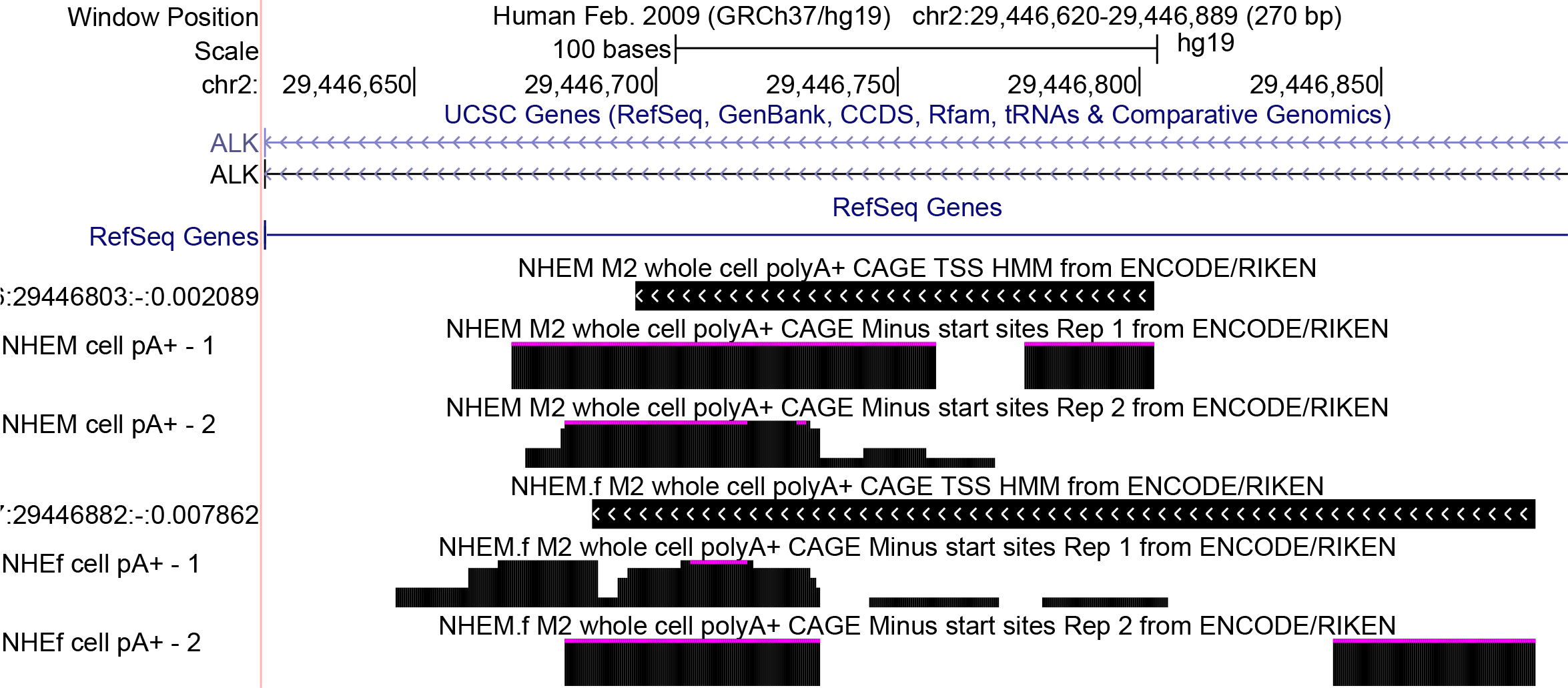
Displayed in the UCSC Genome Browser (http://genome.ucsc.edu) are tracks corresponding to CAGE data for normal human melanocyte cell cultures NHEM_M2 and NHEM.LM2 studied by ENCODE as well as TSSes predicted with Hidden Markov Models from pooled replicates in the ALK gene for *hg19.* Observe that one model predicts a TSS in the region chr2:29,446,803-29,446,696 and the other predicts a TSS in the region chr2:29,446,882-29,446,687, both of which contain the TSS region identified for *ALK^ATI^* in [29], chr2:29,446,768-29,446,744.

